# To nest or not to nest: environmental cues for olive ridley mass nesting events in Odisha, India

**DOI:** 10.1101/2024.10.29.620945

**Authors:** Prahalad Srikanthan, Chandana Pusapati, Akash Verma, Gagan Dhal, Muralidharan Manoharakrishnan, Kartik Shanker

## Abstract

Sea turtles of the genus *Lepidochelys* take part in synchronised nesting events, known as *arribadas*, where a large number of females emerge simultaneously from the sea to nest on the beach. Despite some evidence for the role of environmental factors, there is considerable uncertainty about both the factors that influence mass-nesting at an annual scale as well as the onset of individual events. We used a multi-decadal dataset from two olive ridley mass nesting beaches in Odisha, India, to identify the environmental factors associated with inter-annual nesting variation, nesting seasonality and the onset of arribadas. Interestingly, the two sites showed different patterns in inter-annual patterns associated with the non-occurrence of arribadas, despite being only 250 km apart. The nesting season was consistently during the dry and cooler part of the year. Arribada start dates were non-uniformly distributed across the lunar phase in Gahirmatha with most arribadas occurring during the first and third quarters. Fast Southern Winds characterised the period before and during the arribada at both sites. Machine learning models (XGBoost) identified the most important parameters for the onset of mass nesting and accurately predicted more than 75% of the start dates. Lunar phase, mean sea level pressure and wind speed and direction were important parameters at both sites. Differences in parameter importance between sites suggest that mass nesting is influenced by the interaction of several environmental parameters, along with site-specific characteristics. Insights into the triggers of mass nesting can thus provide guidance for site-specific management efforts.

## Introduction

Reproductive synchrony is a strategy where individuals carry out parts of their reproductive cycle at the same time as other members of the population (Burger 1980; Findlay and Cooke 1982). This strategy is widely observed across the plant and animal kingdoms, such as the simultaneous flowering of bamboo, the emergence of cicadas, or synchronised spawning in several aquatic insects, corals and fish (Corbet 1964; Janzen 1976; Williams and Simon 1995). Studies suggest that breeding or reproductive synchrony serves as an anti-predator strategy and/or increases the chance of finding a mate (Darling 1938; Hatchwell 1991; Descamps 2019). This synchronous behaviour can be mediated by several environmental and biological factors in marine organisms, such as the lunar phase, tidal cycles, temperature, rainfall, and pheromones (Skov et al. 2005; Gerlach 2006; Salminen and Hoikkala 2013; Sen Majumder and Bhadra 2015). For instance, some coral communities in the Great Barrier Reef spawn together in response to changes in temperature, while some corals in the Caribbean use lighting conditions as a cue for gamete release (Clifton 1997; Baird et al. 2009). Similarly, some crabs synchronise their reproduction with lunar/tidal patterns, while precipitation triggers breeding in certain frog species (Skov et al. 2005; Hirschfeld and Rödel 2011). In sea turtles, temperature and rainfall have been shown to influence the breeding phenology in various ways (Pike 2009; Weishampel et al. 2010; Neeman et al. 2015).

Given the close interaction between seasonal changes in environmental parameters and the onset of reproduction, the rapid pace of anthropogenic climate change and its resultant effect on large scale environmental variation can lead to a shift in breeding phenology across various taxa (Parmesan 2006; Yang and Rudolf 2010). Studies have shown that in response to changing environmental conditions, birds have advanced their breeding and migration times, and plants have altered their flowering patterns (Crick et al. 1997; Fitter and Fitter 2002; Møller et al. 2010). In addition to changes in breeding phenology, climatic change can also result in phenological mismatches (Visser et al. 1998). As different climatic cues, such as rainfall and temperature, undergo varying degrees of change, organisms maybe exposed to suboptimal conditions, thereby affecting reproduction and survival (Forrest and Miller-Rushing 2010; Stocker 2014; Iler et al. 2021; Zettlemoyer and DeMarche 2022).

Olive ridley turtles (*Lepidochelys olivacea*) are one of two ridley species which exhibit a prominent synchronous nesting behaviour called the *arribada* (Smith et al. 1980; Plotkin 2007), where a large number of nesting females, ranging from a few thousands to hundreds of thousands, nest simultaneously over a short time period (Plotkin 2007). Although olive ridley turtles participate in solitary nesting as well and are found nesting on tropical coasts all across the world, they are well known for their *arribadas* which occur primarily on a few beaches in Central America and India (Shanker et al. 2021). This behaviour is believed to be a predator satiation strategy, as the nests laid during the arribada experience significantly less predation than solitary nests given their sheer number on the beach (Eckrich and Owens 1995).

Currently, there are a few large rookeries in Mexico, Costa Rica and India that receive over 100,000 nests during a single arribada event and a few smaller mass nesting sites on various coasts across the world, mostly in Pacific Central America (Shanker et al. 2021). However, there is a difference in the frequency and timing of arribada occurrence across these sites. For instance, multiple arribadas occur in Costa Rica throughout the year, but typically only one or two occur in India (Shanker et al 2004). There are currently two mass nesting sites on the east coast of India in Odisha, at Gahirmatha and Rushikulya; at both sites, arribadas take place only during the dry season (Shanker et al. 2004). It is, however, not uncommon for arribada to not occur during certain years at a site, for reasons still unknown.

Despite decades of monitoring arribadas, the triggers of mass nesting are still largely unknown. Previous studies have found that arribadas are triggered by a variety of environmental cues such as wind bursts, salinity of the beach sand, ocean currents, sea surface height, and/or lunar cycles (Plotkin 2007; Barik et al. 2014; Bézy et al. 2020; Poornima 2021). Other studies have theorised that social facilitation or pheromones may act as triggers for an arribada (Mora and Robinson 1982; Owens et al. 1982; Plotkin 2007).

Given the variation in mass nesting events across sites, it is important to study this at a regional scale and determine if the triggers are consistent for populations across and within regions. In this study, we examined the influence of various environmental parameters on arribadas in Gahirmatha and Rushikulya using a ∼40-year dataset. First, we assessed the influence of environmental parameters on the occurrence of arribadas across years (i.e., inter-annual variation). We then examined variation in environmental parameters between the nesting and non-nesting seasons (i.e., intra-annual variation). Finally, we analysed arribada timing (date of onset) at the two mass nesting sites and developed a machine learning model to identify the triggers for arribadas at both these sites.

## Methods

### Study Area

The Rushikulya mass nesting site is located on the southern coast of the Indian state of Odisha, at the mouth of the Rushikulya river (Figure 1). Mass nesting mostly occurs on a 4 km beach stretch that extends north of the river mouth or sometimes on a sand spit that forms in front of the nesting beach (Barik et al. 2014). The mass nesting beach at Rushikulya is highly dynamic due to the seasonal flooding of the river (Chandarana et al. 2017). The Gahirmatha mass nesting beach, located about 250 km north of Rushikulya, is at the mouth of the Brahmani and Baitarani rivers. Nesting used to occur on a 10 km stretch on the mainland beach, until a cyclone in 1989 washed away the nesting beach, creating a 5 km sand spit away from the mainland (Shanker et al. 2004). Later, the sand spit was further divided into smaller islands that change annually (Prusty et al. 2000; Behera and Kaiser 2020).

**1).**
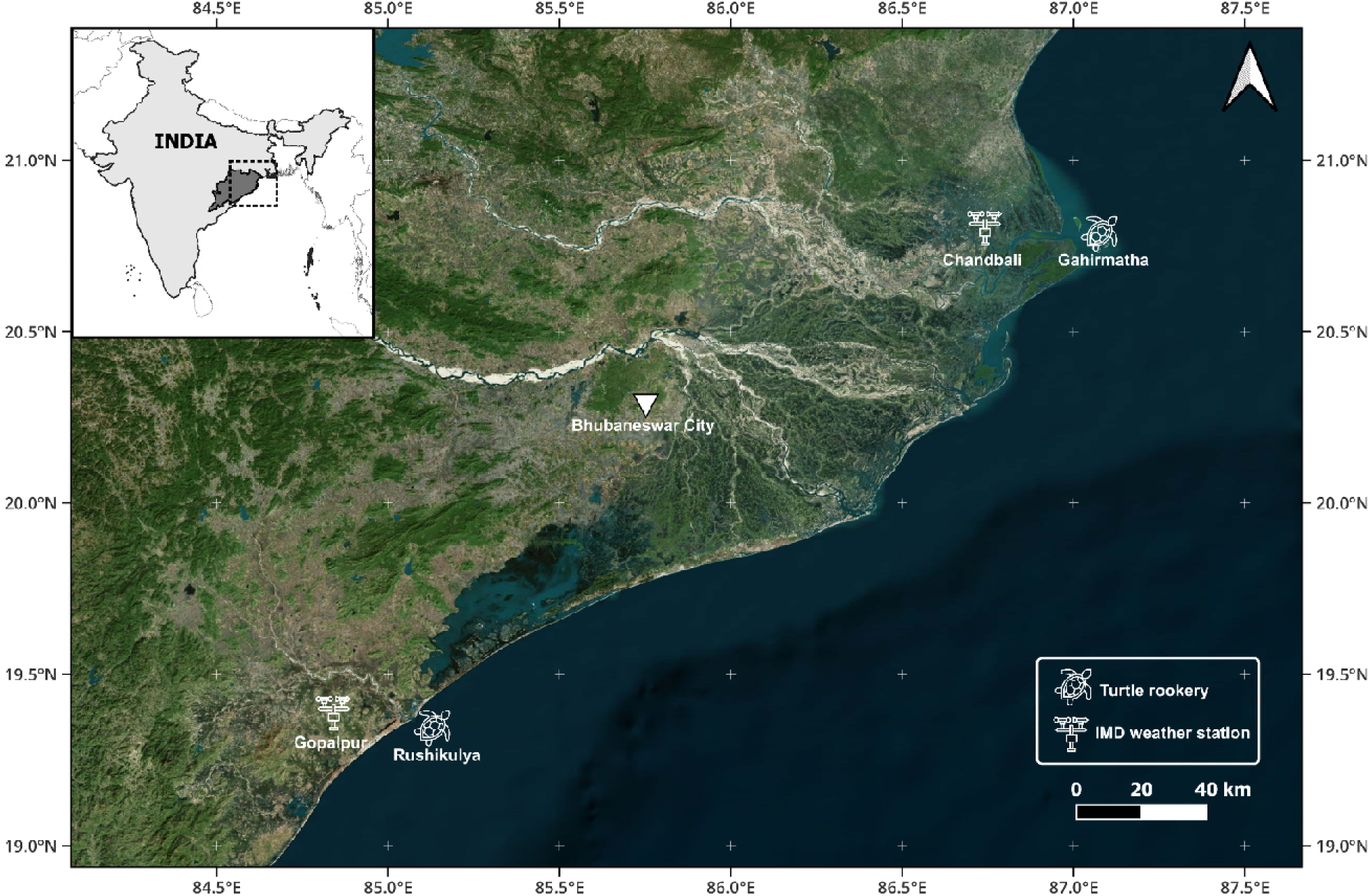
Map of the study area

### Data collection

The arribada occurrence data for Gahirmatha (from 1978 to 2020) was obtained from Shanker et al. (2004) and Poornima (2021); and Rushikulya data (from 1994 to 2020) was obtained from Tripathy (2005), Chandarana et al. (2017) and Shanker et al. (unpubl. data). Meteorological data from 1978 to 2020 was obtained from the Gopalpur weather station, Indian Meteorological Department of the Government of India. This included daily rainfall data, wind speed, wind direction, wet and dry bulb temperature (DBT), average temperature, minimum and maximum temperature, dew point temperature, relative humidity, and Mean Sea Level Pressure (MSLP). Data at two time stamps (17:30 and 08:30) were obtained for the following variables: wind speed, wind direction, DBT, relative humidity, and average temperature. The 17:30 data is the value measured at 17:30, but the 8:30 data is the average value measured from 17:30 the previous day to 08:30. We used the 17:30 data for two reasons: firstly, this time point is more indicative of the general window when arribadas typically commence; secondly, our preliminary analysis indicated lower variation in mean night temperatures across the window that arribadas typically occur. Hence, all the variables used in this study refer to the 17:30 data unless mentioned otherwise.

Our choice of parameters for each analysis was determined based on previous research and available data (Coria Monter 2017; Bézy et al. 2020). Comprehensive environmental data was not available for Chandbali, the weather station closest to Gahirmatha, with the exception of temperature data. However, we observed a high correlation between the temperature data obtained from the Chandbali weather station in Gahirmatha and the Gopalpur weather station in Rushikulya. Thus we utilized the Gopalpur dataset for all environmental parameters.

Monthly indices for the El Nino Southern Oscillation (ENSO) and Indian Ocean Dipole (IOD) were obtained from the NOAA Physical Sciences Laboratory, Boulder Colorado. The ENSO is a cyclical climatic phenomenon that causes abnormally hot (El Niño) or cold (La Niña) conditions in the tropical Pacific. It usually occurs in conjunction with variations in the Southern Oscillation, which is an atmospheric pressure pattern related to Pacific trade winds (McPhaden et al. 2006). The IOD is a similar phenomenon observed in the tropical Indian ocean (Saji et al. 1999). Values of sea surface temperature (SST) were obtained from monthly averages from the HadISST v1.1 dataset (Rayner et al. 2003). Points near Rushikulya and Gahirmatha were sampled to extract SST values using the ‘raster’ package in R (Hijmans 2020). We obtained lunar phase data using the ‘Lunar’ package in R (Lazaridis 2022). To correct any potential discrepancies in the lunar data, we cross verified it with the Islamic calendar, which follows the new moon as the start of each month (Allouche 2002). All statistical analyses were carried out using R (R core team, 2020).

### Statistical Analysis

#### Inter- and intra-annual variation

To understand the influence of various environmental parameters on arribada occurrence for each site, we compared parameters across three groups: years with an arribada (‘Arribada’), years without an arribada (‘No Arribada), and the year that preceded a year without an arribada (‘No_Arribada-1’) using a multinomial logistic regression. The following environmental parameters were used: yearly maximum values of ENSO and IOD, total annual rainfall, yearly average SST, DBT, and MSLP.

To investigate the variation in environmental parameters within years, we compared values between the nesting season (January-April), hatching season (March-May) and the rest of the year (June to December) without separating the sites for the DBT and total rainfall using a multivariate analysis of variance test (MANOVA) (Pillai 2014).

#### Arribada triggers

To identify environmental triggers, we examined the following variables: DBT, wind speed, MSLP, and wind direction. Wind direction was also converted into two continuous circular variables Wind_Direction_EW and Wind_Direction_NS, using a sine and cosine function respectively, to capture the East-West and North-South components.

We first compared these variables across a broader timescale (months to weeks) and then a finer time scale (each of the five days leading up to the arribada) (Table 1a). This allowed us to examine the parameters at both coarse and fine timescales using multinomial logistic regression with the arribada start date as the reference level. In addition, to investigate the possibility of an interaction between wind speed and direction, wind was classified into four categories based on wind speed throughout the nesting season (Table 1b). These were merged to create ‘Wind Class,’ which accounted for both the speed and direction of the winds. The groups were compared across similar coarse and fine timescales as described above.

**Table 1 (a).** Periods used for the fine and coarse scale analysis.

| Variable Name | Type of Data | Data used |
| --- | --- | --- |
| Arri | Single day | Arribada start dates |
| Arri-1 | Single day | 1 day before an arribada start date |
| Arri-2 | Single day | 2 days before an arribada start date |
| Arri-3 | Single day | 3 days before an arribada start date |
| Arri-4 | Single day | 4 days before an arribada start date |
| Arri-5 | Single day | 5 days before an arribada start date |
| Arri-1w | Cumulative | Arri-1 to Arri-7 |
| Arri-2w | Cumulative | Arri-8 to Arri-14 |
| Arri-1M | Cumulative | Arri-14 to Arri-30 |
| Arri-2M (Statistical only) | Cumulative | Arri-31 to Arri-60 |
| Non-nesting Season (Visual only) | Cumulative | May to November |
| Post_Arri | Cumulative | Arri+1 to Arri+30 |
| Nesting Season (Visual only) | Cumulative | December to April |

**Table 1 (b).** Classes for wind speed and direction.

| **Speed (percentile)** | **Class** | **Direction** | **Class** |
| --- | --- | --- | --- |
| 0-25<sup>th</sup> | Slow | E/ENE/ESE/W/WNW/WSW | E/W |
| 25<sup>th</sup>-50<sup>th</sup> | Moderate | S/SE/SW/SSW/SSE | Southern |
| 50<sup>th</sup>-75<sup>th</sup> | Fast | N/NE/NW/NNW/NNE | Northern |
| >75<sup>th</sup> | Very Fast | None | Zero Degrees |

Additionally, we converted wind direction into radians and conducted a Rayleigh test, using the ‘circular’ package in R to assess if wind direction played a role in arribada initiation (Rayleigh 1880; Agostinelli and Lund 2022).

To analyze the effect of lunar phases, we plotted the distribution of the moon phase on the arribada start dates for each site. Similar to wind direction, we conducted a Rayleigh test to investigate if arribadas started uniformly across all phases (Rayleigh 1880; Agostinelli and Lund 2022). This was done using radian values for each phase, obtained from the lunar package, and included data from 1978 to 2023 (Lazaridis 2022). This extended period, compared to the 1978–2020 range used in other analyses, was chosen to incorporate data on the two arribadas that occurred at Rushikulya in 2022 and 2023.

#### Machine learning model

We used eXtreme Gradient Boosting (XGBoost) to create two models, one for each site, to identify the parameters that influence the onset of an arribada. XGBoost was selected due to its ability to accommodate extensive hyperparameter tuning, enabling us to optimize the model’s performance for the specific characteristics of our dataset. This approach proved more effective than traditional models such as random forest and logistic regression.

XGBoost uses a gradient boosting decision tree algorithm that can be used for both classification and regression problems (Chen and Guestrin 2016).

Pairwise correlations were conducted to include only orthogonal parameters in the model. Relative humidity was logit-transformed because the values are presented as percentages. Following Bezy et al. (2020), we converted the lunar phase into two continuous parameters by using cosine and sine transformations. The first parameter ‘Lunar phase’ represented the phase of the moon (1 full, 0 quarter, -1 new moon), whereas ‘Lunar state’ indicated the waxing or waning state of the moon (1 first-quarter, 0 full/new, -1 last-quarter moon.

The response parameters were coded as 1 if an arribada started on that day, and 0 for all other days. Since there are 0, 1 or 2 arribada start dates each year, and all other dates represent 0s (not an arribada start date), the dataset was highly zero-inflated. Thus, up-sampling of the data was conducted to balance the dataset and improving model performance. Synthetic Minority Oversampling Technique (SMOTE) was used to increase the proportion of the minority class from less than 0.2% to 3% for both sites (Chawla et al. 2002). The data was divided into a test (20%) and training (80%) dataset.

The models were created using the ‘xgboost’ R package. We ran 1,500 models for each site and averaged the results to account for randomness in the initial parameters and subsampling within XGBoost. The area under the curve (AUC) of the receiver operating characteristic (ROC) curve and confusion matrices built using the test dataset were utilised to validate the models (Zou et al. 2007; Kuhn 2008; Robin et al. 2011). SHapley Additive exPlanations (SHAP) values were used to determine parameter importance. SHAP values are used to explain the prediction of each data point by computing the contribution of each feature to the prediction. This is done by considering all possible combinations of features, assessing how the model’s prediction changes when each feature is added or removed, and then averaging these contributions to determine the importance of each feature. They are unaffected by tree structure unlike the feature importance values provided by XGBoost (Lundberg and Lee 2017; Lundberg et al. 2018; Liu and Just 2020).

## Results

### Arribada occurrence

Thirty-nine arribadas occurred in Gahirmatha between 1978 and 2020. Arribadas did not occur in 1988, 1992, 1997, 1998, 2002, and 2014 (Shanker et al. 2004; Poornima 2021). There have been 23 arribadas in Rushikulya between its discovery in 1994 and 2020, with arribadas not occurring in 1999, 2000, 2002, 2007, 2016, and 2019 (Pandav et al. 1994; Chandarana et al. 2017; Shanker et al, unpubl. data).

### Inter- and intra-annual patterns

To investigate the effect of variables across years, we compared them across three groups; years with an arribada (‘Arribada’), years without an arribada (‘No_Arribada’), and years preceding a year without an arribada (‘No_Arribada-1’).

Inter-annual patterns: In Gahirmatha, differences between groups were statistically significant for DBT, SST, IOD, and MSLP (Multinomial logistic p<0.05; Supplementary Table 1a).

DBT in the ‘Arribada’ years were the lowest and DBT in ‘No_Arribada-1’ years was lower than the ‘No_Arribada’ years (Multinomial logistic p<0.05, Figure 2a). SST for ‘No_Arribada’ years was higher compared to the other groups (Multinomial logistic p<0.05; Figure 2c). MSLP patterns showed that the ‘No_Arribada’ group had significantly higher values compared to ‘Arribada’ years (Multinomial logistic p<0.05; Supplementary Figure 2c).

**2).**
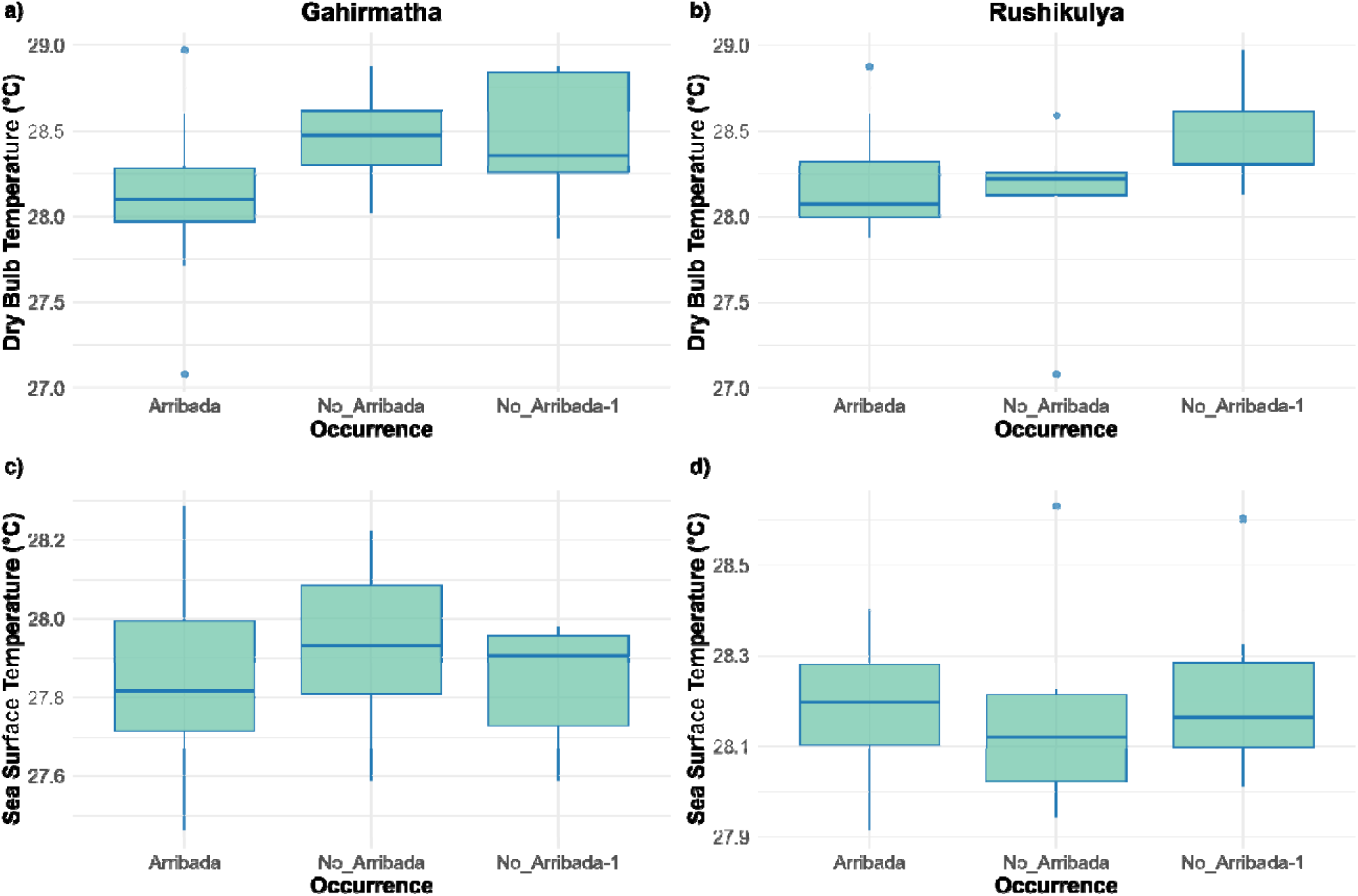
Inter-annual trends. (a-b) Dry bulb temperature (DBT) for years with arribada, no arribada, and years preceding a year without an arribada for Gahirmatha and Rushikulya, respectively. (c-d) Sea surface temperature (SST) across years with arribada, no arribada, and years preceding a year without an arribada for Gahirmatha and Rushikulya, respectively.

In Rushikulya, differences in groups were statistically significant for DBT, MSLP, and SST (Multinomial logistic p<0.05; Supplementary Table 1b). SST in the ‘No_Arribada’ years was lower than in ‘Arribada’ years (Multinomial logistic p<0.05; Figure 2d).

Intra-annual patterns: Total rainfall in the nesting and hatching season was significantly less than the rainfall observed during the rest of the year (MANOVA p<0.05). DBT was significantly lower in the nesting season compared to the other two groups (MANOVA p<0.05).

### Arribada Triggers

#### Environmental triggers

To identify potential environmental triggers, we compared various parameters on the day of arribada and the preceding period at both coarse (weekly) and fine (daily) timescales. At coarse timescales, wind speeds increased marginally before an arribada, but the effect sizes for wind speed were small and not statistically significant (Figure 3, Supplementary Table 5).

**3).**
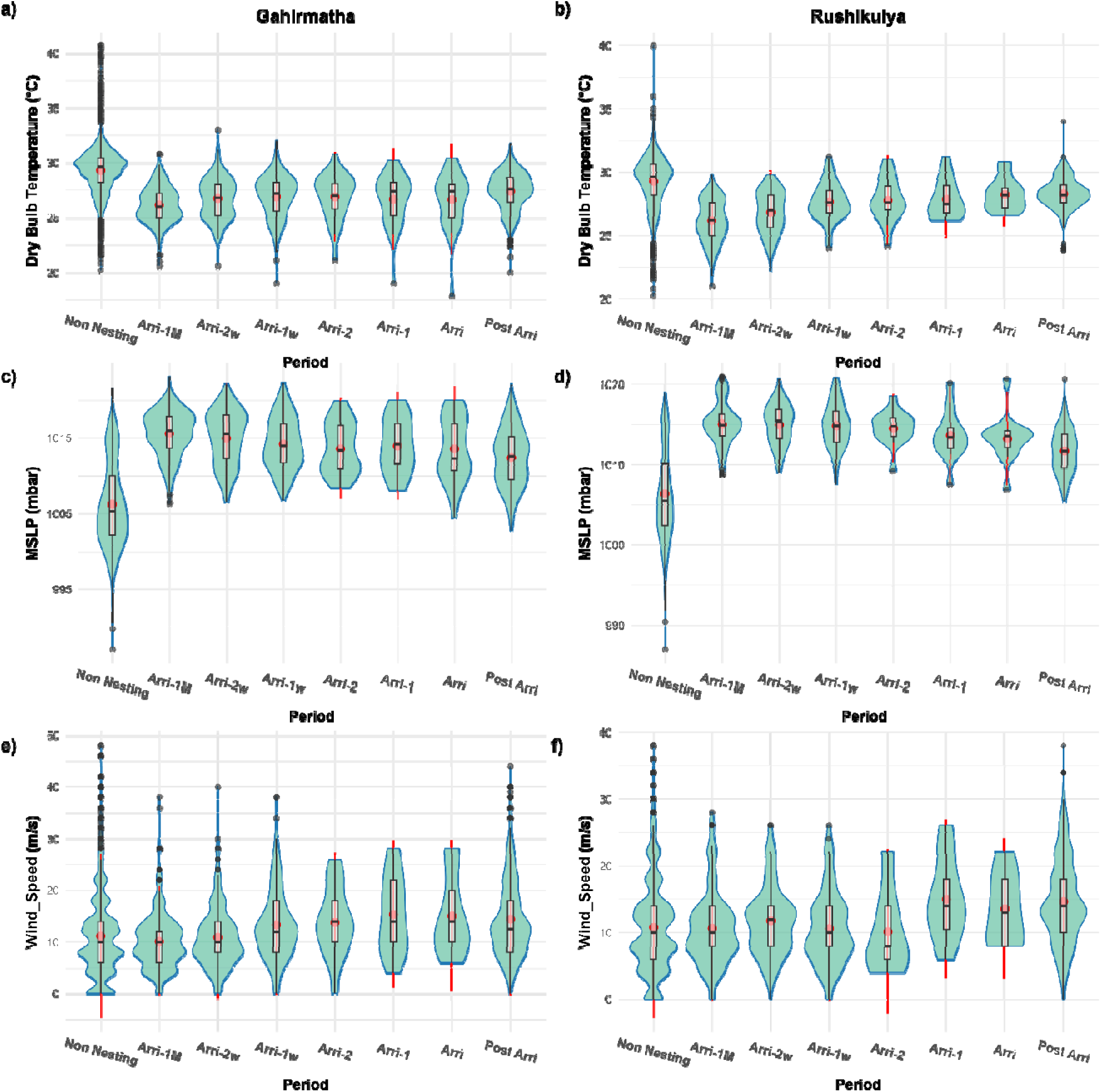
Coarse scale analysis of parameters preceding an arribada. (a-b) Dry Bulb Temperature. (c-d) Mean Sea Level Pressure. (e-f) Wind Speed

At the finer timescale, there was a significantly higher variance in daily temperature one day before the arribada at Gahirmatha (Multinomial logistic p<0.05; Figure 4a). MSLP values were highest one day before the arribada and lowest on the arribada day in Gahirmatha (Multinomial logistic p<0.05; Figure 4c); whereas in Rushikulya, it reached its lowest point one day before arribada with a decreasing trend starting from ‘Arri-5’ onwards (Multinomial logistic p<0.05, Figure 4d).

**4).**
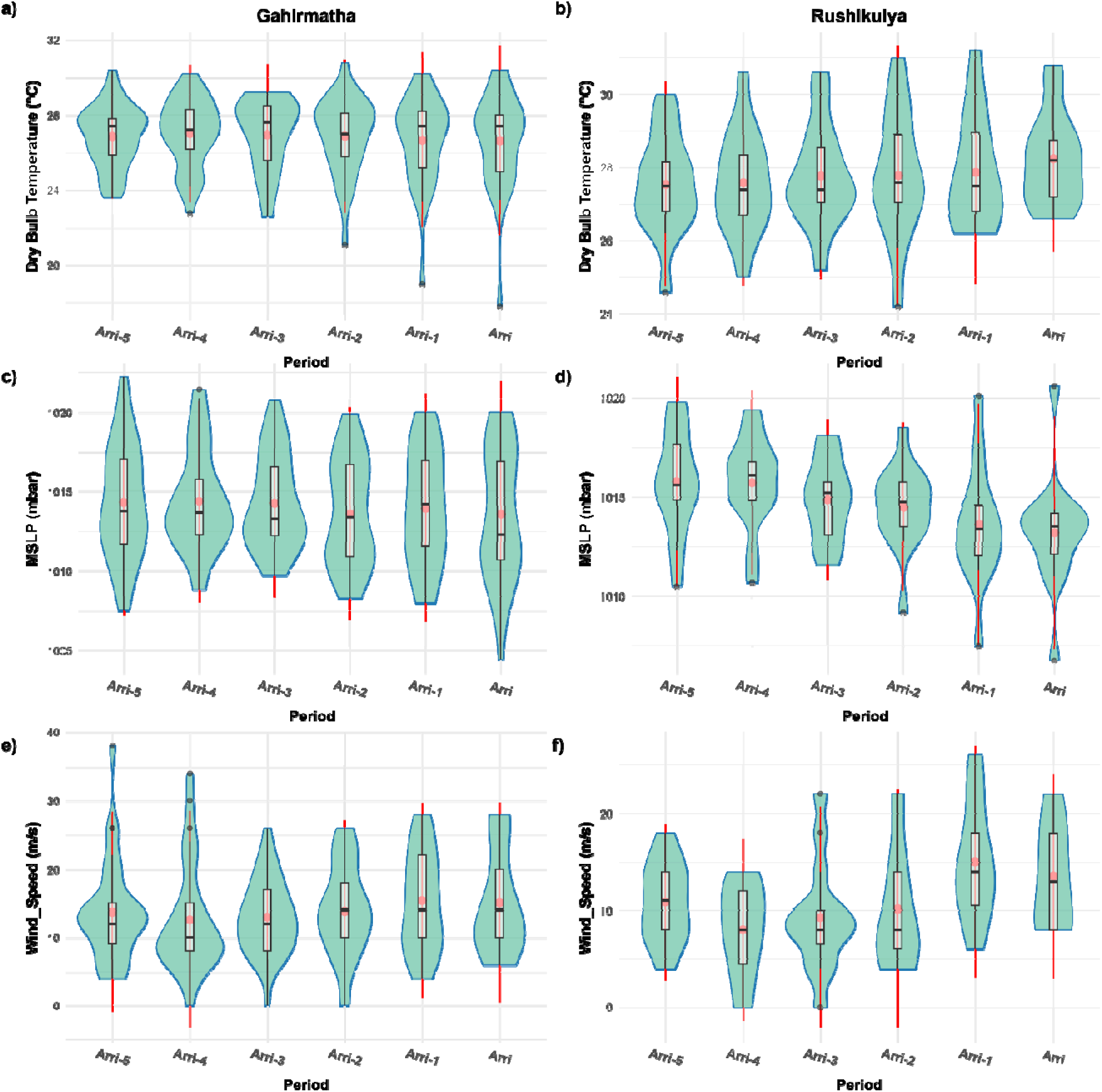
Fine scale analysis of parameters for 5 days preceding an arribada. (a-b) Dry Bulb Temperature. (c-d) Mean Sea Level Pressure. (e-f) Wind Speed

The median wind speed value slightly increased from ‘Arri-3’ to ‘Arri-2’ in Gahirmatha (Figure 4e). A similar pattern, albeit a much larger increase, was observed in Rushikulya from ‘Arri-2’ to ‘Arri-1’ (Figure 4f). On combining wind speed and wind direction, it was observed that southern winds were dominant in the nesting season. Furthermore, ‘Very Fast Southern’ wind was most predominant on two days before an arribada and one day before the arribada in Gahirmatha and Rushikulya respectively, with a gradual increase in frequency from 1 week before the arribada (Figure 5, Supplementary Figure 3). Arribadas occur non-uniformly across wind directions in both sites (Rayleigh test p<0.05). A ‘Very Fast Southern’ wind or a ‘Fast Southern’ wind occurred on over 60% of arribada start dates across sites. In Gahirmatha, ‘Very Fast East’ winds also occurred on an additional 12% of arribada start dates, whereas ‘Moderate South’ or ‘Moderate North winds occurred on 28% and 7% of start dates in Rushikulya.

**5).**
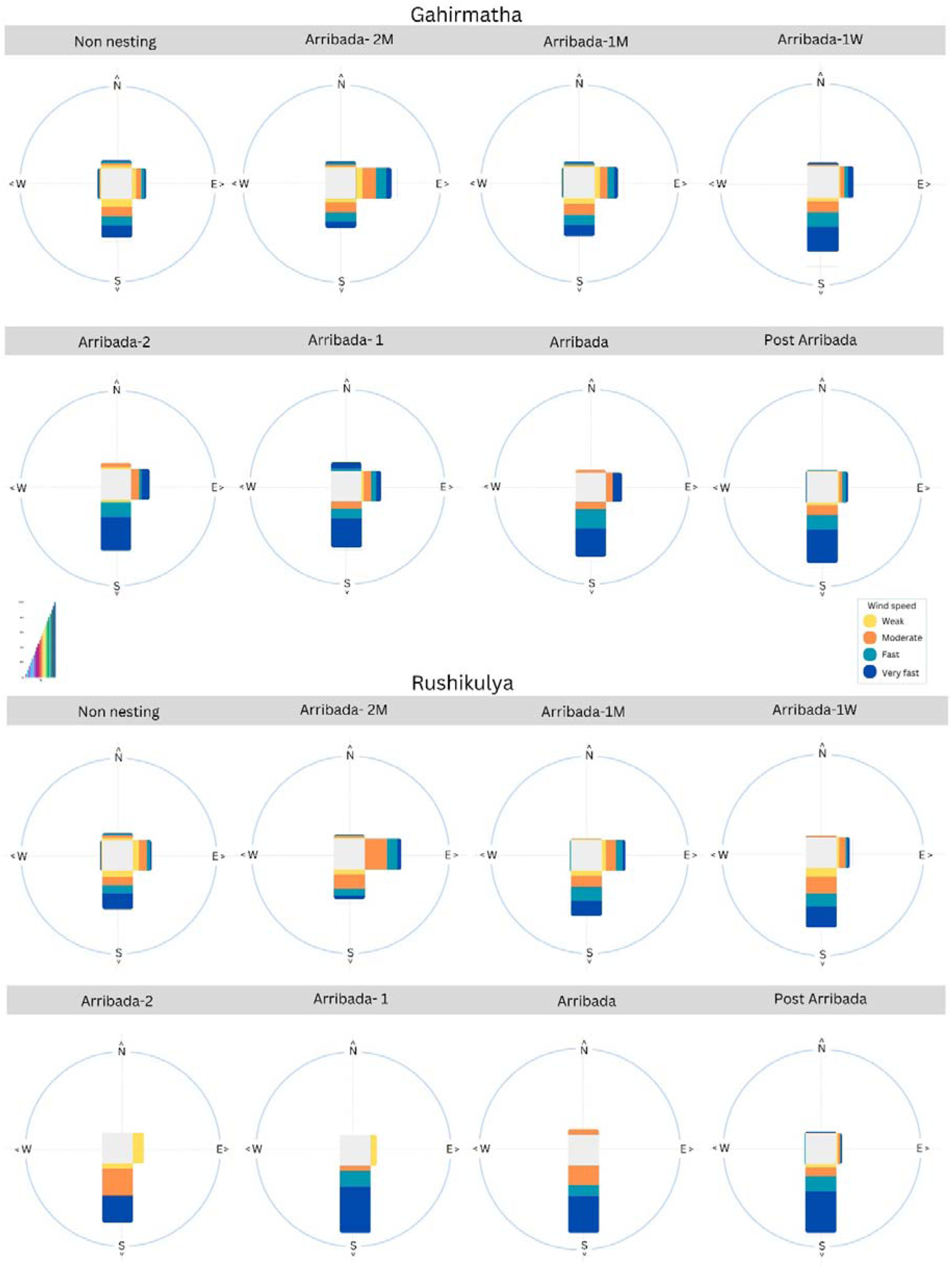
Wind speed and direction preceding an arribada

#### Lunar phase

The analysis of arribada start dates across lunar phases in Gahirmatha revealed that of the 39 arribadas, twelve started in the last quarter, followed by eight during the waning gibbous phase and six in the first quarter (Figure 6). In Rushikulya, of 25 arribadas analysed, eight began during the waxing crescent phase, five during the waning gibbous and four during the last quarter (Figure 6). The Rayleigh test suggested a non-random distribution of start dates across lunar phases for Gahirmatha (Rayleigh test p<0.05), while Rushikulya showed a random distribution (Rayleigh test p=0.17).

**6).**
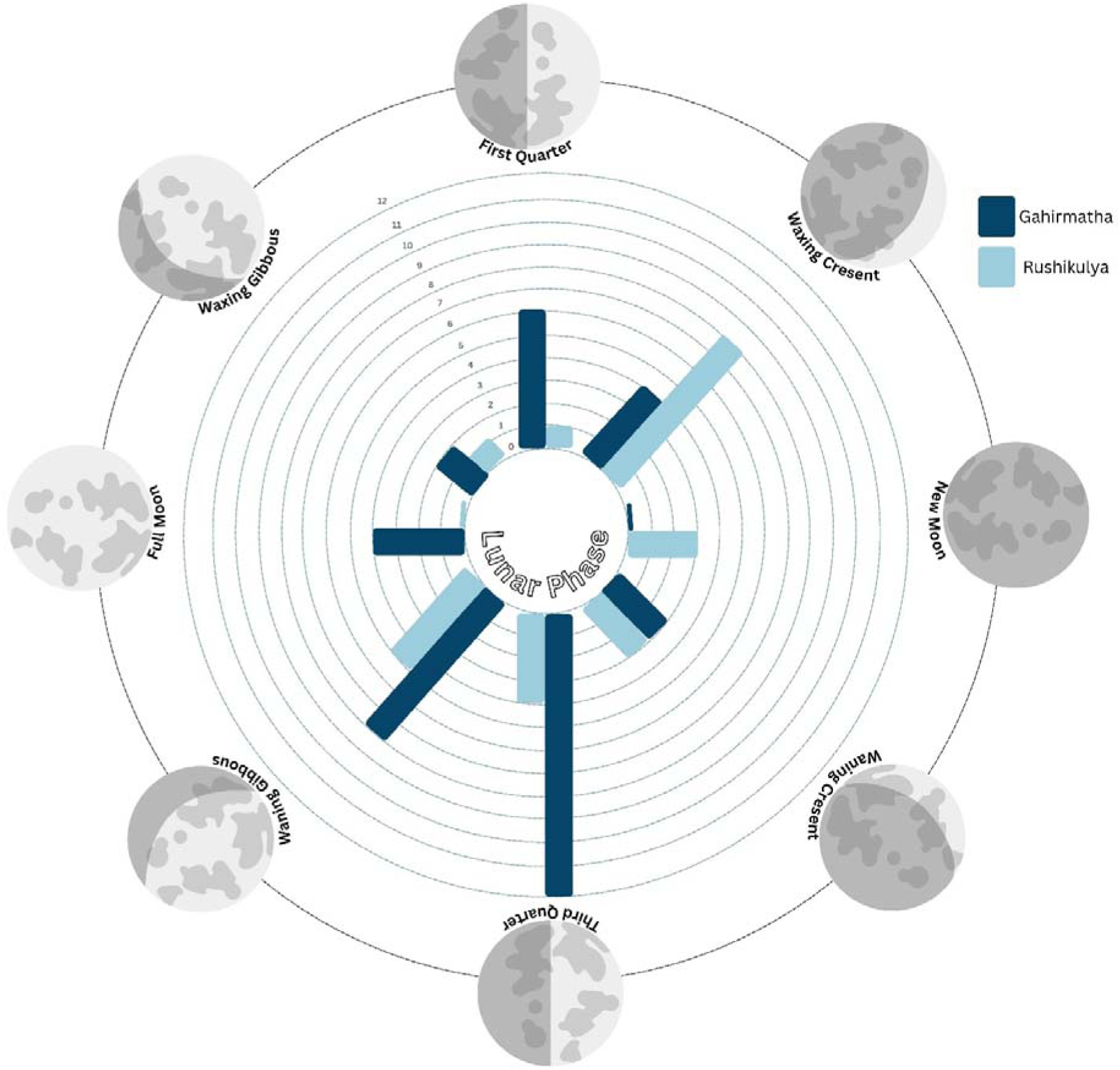
(a-b) Bar plot of lunar phases on arribada start dates for Gahirmatha and Rushikulya, respectively

### Machine Learning Model

To understand which parameters influenced the onset of mass nesting, we developed an XGBoost model for each site. Following up-sampling, the Gahirmatha training set had 4340 zeroes and 149 ones, while the initial test dataset had 1085 zeros and 37 ones. The Rushikulya training set consisted of 3584 zeroes and 101 ones, with the initial test dataset having 896 zeros and 25 ones. The Gahirmatha model had an average AUC of 0.886 and correctly classified 29 out of 37 arribada start dates from the test dataset (Supplementary table 5a, Table 2a). ‘Lunar Phase’, which signifies the phase of the moon, had the highest SHAP value followed by ‘Wind Speed’, ‘Lunar State’, and ‘MSLP’ (Figure 7a). Intermediate values of ‘Lunar Phase’ and lower values of ‘Lunar State’ had a positive impact in classifying arribadas, both of which represent phases closer to the last-quarter moon. Higher wind speeds on the day of the arribada also had a positive impact.

**7).**
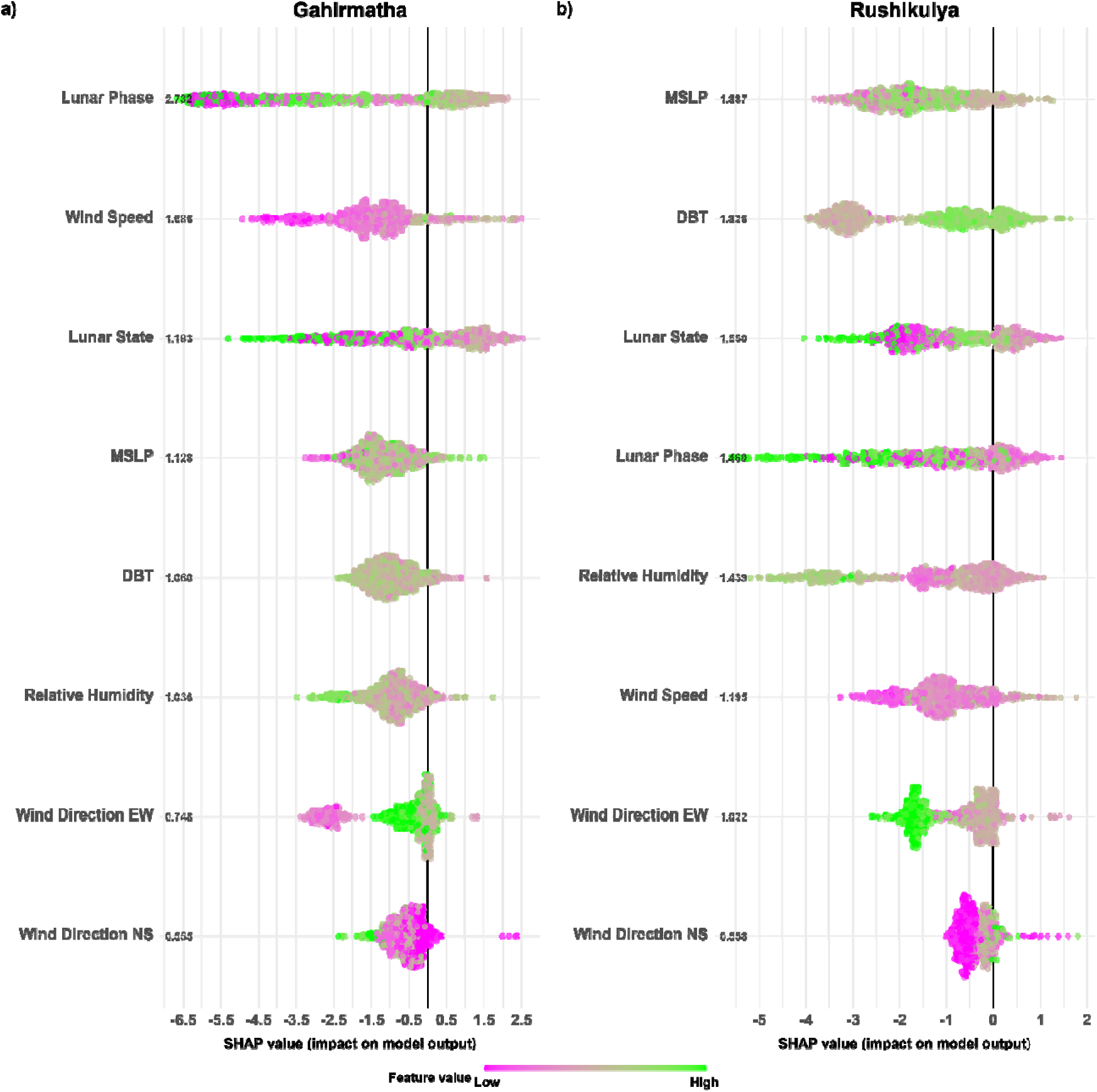
Model output and performance on the test dataset. (a) Parameter SHAP values for onset mass nesting in Gahirmatha. (b) Parameter SHAP values for onset mass nesting in Rushikulya. Each point represents a data point, with colours showing the value of the point. A positive SHAP value indicates a positive impact on classifying an arribada start date.

**Table 2.** Average of Confusion matrices for 1500 XGboost models to evaluate the performance of the classification model by comparing the predicted classes against the actual classes.

a) Gahirmatha
| n= 1122 | **Predicted: 0** | **Predicted: 1** |
| --- | --- | --- |
| **Actual: 0** | **1083** | **2** |
| **Actual: 1** | **8** | **29** |
| Sensitivity = 0.773 | Specificity = 0.998 | Precision = 0.946 |
| Accuracy = 0.991 | F1=0.849 | Kappa = 0.844 |

b) Rushikulya
| n= 921 | **Predicted: 0** | **Predicted: 1** |
| --- | --- | --- |
| **Actual: 0** | **896** | **0** |
| **Actual: 1** | **3** | **22** |
| Sensitivity = 0.898 | Specificity = 0.999 | Precision = 0.986 |
| Accuracy = 0.996 | F1=0.939 | Kappa = 0.937 |

The Rushikulya model had an average AUC of 0.9480 and correctly classified 22 out of 25 start dates in the test dataset (Supplementary Table 5a, Table 2b). MSLP had the highest SHAP value followed by ‘DBT’. The next most important parameters ‘Lunar State’ and ‘Lunar Phase’ had similar SHAP values (Figure 7b). Higher DBT values and intermediate MSLP values on the day of arribada positively influenced the classification of arribadas.

Contrary to Gahirmatha, periods near the new moon positively impacted the model. In addition, lower values of ‘Relative Humidity’ positively impacted the model. (See supplementary Table 5b and 5c, for parameter importance values obtained from XGBoost for Gahirmatha and Rushikulya, respectively).

## Discussion

Since their discovery in the 1970s and 1990s respectively, Gahirmatha and Rushikulya have hosted some of the largest olive ridley arribadas in the world (Bustard 1976; Pandav et al. 1994; Shanker et al. 2004; Shanker et al. 2021). The large number of hatchlings produced from these sites contribute significantly to the stability of the population at a regional level. Therefore, identifying the triggers of mass-nesting is crucial for the conservation and management of this species. In the context of climate change, understanding the climatic preferences of large nesting populations is necessary to devise effective management strategies and mitigate threats.

While the environmental drivers of arribada occurrence at inter-annual scales were equivocal, temperature and rainfall clearly determined seasonality within years. Our results also elucidate the environmental triggers of arribada initiation such as lunar phase, wind speed, and temperature, which have been previously hypothesized to play an important role (Pritchard 1969; Hughes and Richard 1974; Plotkin 2007). Interestingly, our analysis showed that turtles responded slightly differently to cues in Gahirmatha and Rushikulya, despite their being only 250 km apart. This is similar to the dynamics previously observed in the Ostional and Nancite beaches in Costa Rica, which are about 100 km apart (Hughes and Richard 1974; Plotkin 2007). The results suggest that site-specific characteristics interact with environmental parameters to influence the onset of mass nesting.

### Inter- and intra-annual patterns

We compared values across years with an arribada, without an arribada, and years preceding a year without an arribada. Differences in patterns were observed across sites for several parameters. IOD showed a significant effect on arribada occurrence, with a negative phase being associated with the year preceding a year with no arribada. While IOD has not previously been linked to turtle nesting, the higher sea surface temperatures or associated climate phenomena observed on the east coast during a negative IOD phase may disrupt nesting patterns. ENSO showed no effect on arribada occurrence, although ENSO has been previously shown to affect the nesting of green turtles (Limpus and Nicholls 2000; Foley et al. 2006).

SST patterns differed among sites, with the values for years without an arribada being higher than for years with an arribada in Gahirmatha, whereas the opposite occurs in Rushikulya. Despite these differences in SST patterns, the large effect sizes observed show that years without an arribada consistently have higher SSTs at both Gahirmatha and Rushikulya (Supplementary Table 1b). Similarly, rainfall was lower in years without an arribada in Rushikulya, but no such pattern was seen in Gahirmatha. These contradictory patterns imply that large scale environmental factors may interact with local beach dynamics to play a role in the occurrence or non-occurrence of arribadas which needs further elucidation. At both sites, beach erosion due to the proximity to the river mouth and periodic cyclones has resulted in significant variation beach width and location (Barik et al. 2023; Mishra et al. 2024). Thus, the availability of suitable nesting beach may have a significant influence on the occurrence of arribadas in that year.

To identify if turtles prefer certain environmental conditions for nesting or hatching, we compared parameters across the nesting, hatching, and non-nesting seasons. We observed similar patterns for temperature and rainfall between sites with significantly lower values for both parameters during the nesting season. The temperature range during winter may offer the best opportunity to produce both male and female hatchlings. Arribadas at Rushikulya produced slightly female biased clutches between 2009 and 2020 (Chandarana et al. 2017). Beyond March, temperatures in Odisha are extremely high followed by two consecutive monsoons with a relatively short dry period in between. The Northeast monsoon from October to December often includes severe cyclonic storms which result in inundation and erosion of nesting beaches. Thus, the dry period after this may offer the best opportunity for nesting and hatching success. In fact, olive ridleys on the west coast as well as olive ridley and leatherback turtles in the Andaman and Nicobar Islands follow a similar seasonality (Swaminathan et al. 2017).

### Arribada triggers

To understand which environmental parameters could trigger mass nesting, we compared values across two different timescales; daily and weekly. Additionally, we created a model to identify the parameters that were most influential for the onset of mass nesting.

Turtles are known to react to environmental thresholds or temperature ranges, as observed in freshwater turtles (Bowen et al. 2005). Additionally, air temperatures can affect sea surface temperatures or sand temperatures, which are known to affect nesting in sea turtles (Mazaris et al. 2008; Pike 2008; Laloë et al. 2014). Our analysis shows that temperature is an important trigger for arribadas in Rushikulya. However, we did not observe a similar pattern in Gahirmatha indicating that triggers might be a result of interaction between regional and local site-specific factors. At both sites, the variation in MSLP between the day before the arribada and the arribada start date suggests that olive ridleys may react to pressure differences, similar to loggerhead and leatherback turtles (Pike 2008; Schofield et al. 2010; Palomino-González et al. 2020). Interestingly, there are contrasting patterns across sites where a drop and rise in MSLP act as triggers in Gahirmatha and Rushikulya respectively.

Wind bursts have been associated with the onset of nesting in both anecdotal accounts (Pritchard 1969; Plotkin 2007) as well as in recent studies (Barik et al. 2014). In our analysis, a wind burst was observed one day before the arribada in Rushikulya. A strong southern wind was observed a day before the arribada for more than 60% of the arribadas across sites, with strong winds being observed on over 75% of start dates in Gahirmatha. Strong winds can act as a signal due to their effect on currents (Mohan et al. 2005) and in addition, an increase in wind speed is known to be correlated with an increase in wave height, which could further facilitate the movement of turtles towards the shore (Young and Ribal 2019). Moreover, our analysis supports anecdotal evidence suggesting that southern winds play a significant role as triggers of arribada initiation.

Both the lunar phase and state were shown to be important cues for arribadas at both sites. Most arribadas occur close to the last quarter moon in Gahirmatha, whereas arribadas occur near the waxing crescent phase in Rushikulya. The preference for the quarter moons is thought to be related to predator avoidance or increased prey abundance (Pinou et al. 2009; Witt 2013). Since they are amongst the smallest species of sea turtles, olive ridleys may prefer phases of the moon associated with neap tides. However, we were unable to integrate tidal data into the study due to the lack of daily data that coincides with the timeline of the dataset.

These results provide evidence for the potential triggers of an arribada and the differences observed across sites. Despite the multi-year dataset and the numerous parameters taken into account, our study had certain limitations. Arribada start dates present a small sample size relative to non-start dates, and synthetic data points had to be created to account for the imbalance in the dataset, which could affect the results. In addition, the decision to denote the first day of the arribada is subjective and based on field researchers present at the nesting site. Future studies could also incorporate tidal and oceanographic data on a daily scale or examine the possibility of social signals as triggers for an arribada.

## Conclusion

Through the analysis of environmental factors and machine learning models, this study provides valuable insights into the triggers of mass nesting. By examining triggers across varying timescales, the study highlights the differences in triggers across sites and the importance of wind speed, temperature, and lunar phase in the initiation of an arribada. These findings not only add credence to anecdotal observations but also enhance our understanding of the underlying mechanisms driving mass nesting behaviour. Furthermore, this study provides a practical framework that can be replicated in future research to address skewed datasets. The differences in triggers across sites emphasizes the need for site-specific considerations in conservation efforts.

## Acknowledgements

We would like to thank the Odisha Forest Department for their continued support in carrying out the long-term monitoring project. We are grateful to the frontline staff of the Forest Department that have contributed to the arribada occurrence data at Gahirmatha and Rushikulya. We thank the Dakshin field staff, Madhusudhan Behera, Bipro Behera, Mahendra Nayak, Surendra Behera, Shankar Rao, Magata Behera and Judhistir Behera for their invaluable contribution to the data and the project over the years. We would also like to thank the researchers at Dakshin Foundation, Anusha Koushik, Divya Karnad, Adhith Swaminathan, Ema Fatima, Amrit Kumar Mishra, Nupur Kale, Chetan Rao, Alissa Barnes, Hariprasath R, Avik Banerjee and Mohit Mudliar who contributed to the data through the monitoring project. We would also like to thank Vidisha for her help with the figures.

