## Supplementary for "To nest or not to nest: environmental cues for olive ridley mass nesting events in Odisha, India"

**Table of Contents:**

| **Supplementary Tables** | Page 2 |
| --- | --- |
| **Supplementary Figures** | Page 9 |

*Table S1a.* Inter-Annual Multinomial logistic regression results for Gahirmatha and (b) Rushikulya. Effect size is the log odds ratio, which is e^(estimate). Reference level is ‘No Arribada’

| **Level** | **Variable** | **estimate** | **std.error** | **statistic** | **p.value** | **conf.low** | **conf.high** |
| --- | --- | --- | --- | --- | --- | --- | --- |
| Arribada | (Intercept) | 0.059 | <0.001 | 228.58 | 0.00 | 0.058 | 0.059 |
| Arribada | ENSO | 0.097 | 0.376 | 0.258 | 0.796 | -0.640 | 0.835 |
| Arribada | IOD | -1.46 | 0.970 | -1.51 | 0.132 | -3.36 | 0.440 |
| Arribada | SST | -2.62 | 0.835 | -3.14 | 0.002 | -4.26 | -0.985 |
| Arribada | Rainfall.mm. | 7.8E-4 | 0.002 | -0.506 | 0.613 | -0.004 | 0.002 |
| Arribada | DBT | -1.75 | 0.891 | -1.96 | 0.049 | -3.49 | <0.001 |
| Arribada | MSLP | 0.124 | 0.045 | 2.76 | 0.006 | 0.036 | 0.213 |
| No_Arribada-1 | (Intercept) | -0.024 | <0.001 | -68.45 | 0.00 | -0.025 | -0.023 |
| No_Arribada-1 | ENSO | -0.302 | 0.464 | -0.650 | 0.515 | -1.21 | 0.607 |
| No_Arribada-1 | IOD | -2.31 | 1.12 | -2.06 | 0.039 | -4.50 | -0.114 |
| No_Arribada-1 | SST | -5.03 | 0.586 | -8.59 | <0.001 | -6.18 | -3.89 |
| No_Arribada-1 | Rainfall.mm. | 6.06E-05 | 0.002 | 0.033 | 0.974 | -0.004 | 0.004 |
| No_Arribada-1 | DBT | 3.75 | 0.483 | 7.75 | <0.001 | 2.80 | 4.69 |
| No_Arribada-1 | MSLP | 0.035 | 0.029 | 1.20 | 0.230 | -0.022 | 0.091 |

*Table S1b.* Inter-Annual Multinomial logistic regression results for Rushikulya. Effect size is the log odds ratio, which is e^(estimate). Reference level is ‘No Arribada’

| **Level** | **Variable** | **estimate** | **std.error** | **statistic** | **p.value** | **conf.low** | **conf.high** |
| --- | --- | --- | --- | --- | --- | --- | --- |
| Arribada | (Intercept) | -49.75 | 0.002 | -28441.68 | 0.00 | -49.76 | -49.75 |
| Arribada | ENSO | 0.204 | 0.434 | 0.471 | 0.638 | -0.646 | 1.05 |
| Arribada | IOD | -0.356 | 1.06 | -0.337 | 0.736 | -2.43 | 1.72 |
| Arribada | SST | -1.61 | 0.484 | -3.33 | <0.001 | -2.56 | -0.665 |
| Arribada | Rainfall.mm. | 0.002 | 0.001 | 1.60 | 0.110 | <0.001 | 0.005 |
| Arribada | DBT | 0.504 | 1.33 | 0.380 | 0.704 | -2.10 | 3.11 |
| Arribada | MSLP | 0.079 | 0.047 | 1.70 | 0.090 | -0.012 | 0.171 |
| No_Arribada-1 | (Intercept) | 43.61 | <0.001 | 45932.51 | 0.00 | 43.61 | 43.61 |
| No_Arribada-1 | ENSO | 0.078 | 0.496 | 0.157 | 0.875 | -0.895 | 1.05 |
| No_Arribada-1 | IOD | 1.51 | 1.31 | 1.16 | 0.247 | -1.05 | 4.07 |
| No_Arribada-1 | SST | 0.799 | 0.550 | 1.45 | 0.146 | -0.279 | 1.88 |
| No_Arribada-1 | Rainfall.mm. | 0.001 | 0.002 | 0.754 | 0.451 | -0.002 | 0.005 |
| No_Arribada-1 | DBT | 2.27 | 0.629 | 3.61 | <0.001 | 1.04 | 3.51 |
| No_Arribada-1 | MSLP | -0.131 | 0.030 | -4.32 | <0.001 | -0.191 | -0.072 |

*Table S2.* MANOVA results for the intra-annual analysis

| **Df** | **Pillai** | **approx F** | **num Df** | **den Df** | **Pr(>F)** |
| --- | --- | --- | --- | --- | --- |
| 2 | 0.764756 | 30.95566 | 4 | 200 | 4.63E-20 |
| 100 | NA | NA | NA | NA | NA |

*Table S3a.* Multinomial logistic regression results for 5 days preceding an arribada for Gahirmatha

| **y.level** | **term** | **estimate** | **std.error** | **statistic** | **p.value** | **conf.low** | **conf.high** |
| --- | --- | --- | --- | --- | --- | --- | --- |
| Arri-1 | (Intercept) | -163.07 | <0.001 | -218593 | 0 | -163.07 | -163.07 |
| Arri-1 | DBT | 0.379 | 0.156 | 2.418 | 0.015 | 0.071 | 0.686 |
| Arri-1 | `MSLP (mbar)` | 0.151 | 0.004 | 36.87 | <0.001 | 0.143 | 0.159 |
| Arri-1 | `Wind_Speed (m/s)` | 0.032 | 0.036 | 0.880 | 0.378 | -0.039 | 0.105 |
| Arri-1 | Wind_Direction_EW | -1.03 | 0.492 | -2.10 | 0.034 | -2.00 | -0.073 |
| Arri-1 | Wind_Direction_NS | 2.11 | 0.452 | 4.68 | <0.001 | 1.23 | 3.00 |
| Arri-2 | (Intercept) | -62.56 | <0.001 | -78744.9 | 0 | -62.56 | -62.55 |
| Arri-2 | DBT | 0.25 | 0.155 | 1.65 | 0.098 | -0.048 | 0.562 |
| Arri-2 | `MSLP (mbar)` | 0.056 | 0.004 | 13.77 | <0.001 | 0.048 | 0.064 |
| Arri-2 | `Wind_Speed (m/s)` | -0.018 | 0.037 | -0.488 | 0.625 | -0.092 | 0.055 |
| Arri-2 | Wind_Direction_EW | -0.916 | 0.514 | -1.78 | 0.074 | -1.923 | 0.091 |
| Arri-2 | Wind_Direction_NS | 1.55 | 0.469 | 3.31 | <0.001 | 0.636 | 2.47 |
| Arri-3 | (Intercept) | -147.45 | 0.001 | -124644 | 0 | -147.46 | -147.45 |
| Arri-3 | DBT | 0.243 | 0.162 | 1.50 | 0.133 | -0.074 | 0.562 |
| Arri-3 | `MSLP (mbar)` | 0.140 | 0.004 | 33.20 | <0.001 | 0.131 | 0.148 |
| Arri-3 | `Wind_Speed (m/s)` | -0.030 | 0.038 | -0.803 | 0.421 | -0.106 | 0.044 |
| Arri-3 | Wind_Direction_EW | -0.936 | 0.554 | -1.68 | 0.091 | -2.02 | 0.150 |
| Arri-3 | Wind_Direction_NS | 0.774 | 0.554 | 1.39 | 0.162 | -0.312 | 1.86 |
| Arri-4 | (Intercept) | -167.59 | 0.001 | -164033 | 0 | -167.60 | -167.59 |
| Arri-4 | DBT | 0.248 | 0.162 | 1.53 | 0.125 | -0.069 | 0.566 |
| Arri-4 | `MSLP (mbar)` | 0.160 | 0.004 | 38.13 | 0 | 0.152 | 0.168 |
| Arri-4 | `Wind_Speed (m/s)` | -0.046 | 0.039 | -1.17 | 0.238 | -0.123 | 0.030 |
| Arri-4 | Wind_Direction_EW | -1.51 | 0.546 | -2.77 | 0.005 | -2.58 | -0.446 |
| Arri-4 | Wind_Direction_NS | 0.899 | 0.535 | 1.68 | 0.092 | -0.149 | 1.94 |
| Arri-5 | (Intercept) | -149.44 | 0.001 | -101278 | 0 | -149.44 | -149.44 |
| Arri-5 | DBT | 0.158 | 0.164 | 0.962 | 0.335 | -0.164 | 0.481 |
| Arri-5 | `MSLP (mbar)` | 0.143 | 0.004 | 33.87 | <0.001 | 0.135 | 0.152 |
| Arri-5 | `Wind_Speed (m/s)` | -0.014 | 0.037 | -0.384 | 0.700 | -0.088 | 0.059 |
| Arri-5 | Wind_Direction_EW | -0.759 | 0.578 | -1.31 | 0.189 | -1.89 | 0.374 |
| Arri-5 | Wind_Direction_NS | 0.211 | 0.635 | 0.333 | 0.738 | -1.03 | 1.45 |

*Table S3b.* Multinomial logistic regression results for 5 days preceding an arribada for Rushikulya

| **y.level** | **term** | **estimate** | **std.error** | **statistic** | **p.value** | **conf.low** | **conf.high** |
| --- | --- | --- | --- | --- | --- | --- | --- |
| Arri-1 | (Intercept) | -31.53 | 0.002 | -13266.40 | 0.00 | -31.54 | -31.53 |
| Arri-1 | DBT | -0.103 | 0.283 | -0.364 | 0.716 | -0.658 | 0.452 |
| Arri-1 | `MSLP (mbar)` | 0.032 | 0.008 | 4.14 | <0.001 | 0.017 | 0.049 |
| Arri-1 | `Wind_Speed (m/s)` | 0.058 | 0.070 | 0.833 | 0.405 | -0.079 | 0.195 |
| Arri-1 | Wind_Direction_EW | 0.924 | 1.12 | 0.828 | 0.408 | -1.26 | 3.11 |
| Arri-1 | Wind_Direction_NS | -0.242 | 1.10 | -0.220 | 0.826 | -2.39 | 1.91 |
| Arri-2 | (Intercept) | 35.12 | 0.002 | 17515.91 | 0.00 | 35.11 | 35.12 |
| Arri-2 | DBT | -0.142 | 0.280 | -0.506 | 0.613 | -0.690 | 0.406 |
| Arri-2 | `MSLP (mbar)` | -0.031 | 0.008 | -3.83 | <0.001 | -0.046 | -0.015 |
| Arri-2 | `Wind_Speed (m/s)` | -0.121 | 0.075 | -1.62 | 0.105 | -0.267 | 0.025 |
| Arri-2 | Wind_Direction_EW | 1.21 | 1.23 | 0.987 | 0.324 | -1.20 | 3.62 |
| Arri-2 | Wind_Direction_NS | -1.59 | 1.54 | -1.04 | 0.299 | -4.60 | 1.41 |
| Arri-3 | (Intercept) | -107.77 | 0.003 | -32599.10 | 0.00 | -107.78 | -107.76 |
| Arri-3 | DBT | -0.198 | 0.282 | -0.703 | 0.482 | -0.750 | 0.354 |
| Arri-3 | `MSLP (mbar)` | 0.113 | 0.008 | 14.23 | <0.001 | 0.098 | 0.129 |
| Arri-3 | `Wind_Speed (m/s)` | -0.137 | 0.080 | -1.70 | 0.089 | -0.294 | 0.021 |
| Arri-3 | Wind_Direction_EW | 0.118 | 1.08 | 0.109 | 0.913 | -2.00 | 2.23 |
| Arri-3 | Wind_Direction_NS | 0.082 | 0.822 | 0.099 | 0.921 | -1.53 | 1.69 |
| Arri-4 | (Intercept) | -195.02 | 0.002 | -114862.00 | 0.00 | -195.03 | -195.02 |
| Arri-4 | DBT | -0.169 | 0.293 | -0.575 | 0.565 | -0.743 | 0.406 |
| Arri-4 | `MSLP (mbar)` | 0.196 | 0.008 | 24.09 | <0.001 | 0.180 | 0.212 |
| Arri-4 | `Wind_Speed (m/s)` | -0.168 | 0.087 | -1.94 | 0.053 | -0.338 | 0.002 |
| Arri-4 | Wind_Direction_EW | 2.32 | 0.923 | 2.52 | 0.012 | 0.516 | 4.13 |
| Arri-4 | Wind_Direction_NS | -2.46 | 1.08 | -2.28 | 0.023 | -4.58 | -0.341 |
| Arri-5 | (Intercept) | -205.79 | <0.001 | -221511.00 | 0.00 | -205.79 | -205.79 |
| Arri-5 | DBT | -0.179 | 0.299 | -0.600 | 0.549 | -0.765 | 0.407 |
| Arri-5 | `MSLP (mbar)` | 0.207 | 0.008 | 24.47 | <0.001 | 0.191 | 0.224 |
| Arri-5 | `Wind_Speed (m/s)` | -0.029 | 0.079 | -0.369 | 0.712 | -0.184 | 0.126 |
| Arri-5 | Wind_Direction_EW | 1.88 | 1.21 | 1.55 | 0.121 | -0.496 | 4.25 |
| Arri-5 | Wind_Direction_NS | -0.788 | 1.32 | -0.596 | 0.551 | -3.38 | 1.80 |

*Table S3c.* Effect sizes for parameters one day before an arribada. Effect size is the log odds ratio, which is e^(Coefficient)

| **Site** | **Variable** | **Coefficient** | **Odds Ratio** | **p-value** |
| --- | --- | --- | --- | --- |
| Gahirmatha | (Intercept) | -163.07 | - | 0 |
| Gahirmatha | DBT (C) | 0.379 | 1.46 | 0.015 |
| Gahirmatha | MSLP (mbar) | 0.151 | 1.16 | <0.001 |
| Gahirmatha | Wind_Speed (m/s) | 0.032 | 1.03 | 0.378 |
| Gahirmatha | Wind_Direction_EW | -1.03 | 0.355 | 0.0348 |
| Gahirmatha | Wind_Direction_NS | 2.11 | 8.35 | <0.001 |
| Rushikulya | (Intercept) | -31.53 | - | 0 |
| Rushikulya | DBT (C) | -0.103 | 0.902 | 0.715 |
| Rushikulya | MSLP (mbar) | 0.032 | 1.03 | <0.001 |
| Rushikulya | Wind_Speed (m/s) | 0.058 | 1.06 | 0.404 |
| Rushikulya | Wind_Direction_EW | 0.923 | 2.51 | 0.407 |
| Rushikulya | Wind_Direction_NS | -0.24 | 0.785 | 0.825 |

*Table S4a.* Multinomial logistic regression results for the coarse analysis – Gahirmatha. Effect size is the log odds ratio, which is e^(estimate)

| **y.level** | **term** | **estimate** | **std.error** | **statistic** | **p.value** | **conf.low** | **conf.high** |
| --- | --- | --- | --- | --- | --- | --- | --- |
| Arri-1w | (Intercept) | -30.00 | <0.001 | -180026.00 | 0.00 | -30.00 | -30.00 |
| Arri-1w | DBT | 0.115 | 0.111 | 1.04 | 0.299 | -0.102 | 0.332 |
| Arri-1w | `MSLP (mbar)` | 0.029 | 0.003 | 10.10 | <0.001 | 0.024 | 0.035 |
| Arri-1w | `Wind_Speed (m/s)` | -0.021 | 0.017 | -1.25 | 0.210 | -0.055 | 0.012 |
| Arri-1w | Wind_Direction_EW | -0.778 | 0.146 | -5.33 | <0.001 | -1.06 | -0.492 |
| Arri-1w | Wind_Direction_NS | 0.697 | 0.148 | 4.70 | <0.001 | 0.407 | 0.988 |
| Arri-2w | (Intercept) | -58.18 | <0.001 | -337431.00 | 0.00 | -58.18 | -58.18 |
| Arri-2w | DBT | 0.131 | 0.110 | 1.19 | 0.234 | -0.085 | 0.347 |
| Arri-2w | `MSLP (mbar)` | 0.057 | 0.003 | 19.61 | <0.001 | 0.051 | 0.063 |
| Arri-2w | `Wind_Speed (m/s)` | -0.066 | 0.019 | -3.39 | <0.001 | -0.104 | -0.028 |
| Arri-2w | Wind_Direction_EW | -0.323 | 0.164 | -1.97 | 0.049 | -0.644 | -0.002 |
| Arri-2w | Wind_Direction_NS | 0.273 | 0.169 | 1.61 | 0.106 | -0.058 | 0.605 |
| Arri-1M | (Intercept) | -21.75 | <0.001 | -142561.00 | 0.00 | -21.75 | -21.75 |
| Arri-1M | DBT | -0.034 | 0.106 | -0.321 | 0.748 | -0.242 | 0.174 |
| Arri-1M | `MSLP (mbar)` | 0.027 | 0.003 | 9.56 | <0.001 | 0.021 | 0.032 |
| Arri-1M | `Wind_Speed (m/s)` | -0.082 | 0.017 | -4.71 | <0.001 | -0.116 | -0.048 |
| Arri-1M | Wind_Direction_EW | -0.375 | 0.109 | -3.43 | <0.001 | -0.589 | -0.160 |
| Arri-1M | Wind_Direction_NS | 0.803 | 0.104 | 7.75 | <0.001 | 0.600 | 1.01 |
| Arri-2M | (Intercept) | -48.42 | <0.001 | -348518.00 | 0.00 | -48.42 | -48.42 |
| Arri-2M | DBT | -0.073 | 0.105 | -0.701 | 0.484 | -0.279 | 0.132 |
| Arri-2M | `MSLP (mbar)` | 0.054 | 0.003 | 19.76 | <0.001 | 0.049 | 0.060 |
| Arri-2M | `Wind_Speed (m/s)` | -0.077 | 0.017 | -4.55 | <0.001 | -0.110 | -0.044 |
| Arri-2M | Wind_Direction_EW | -0.070 | 0.095 | -0.731 | 0.465 | -0.257 | 0.117 |
| Arri-2M | Wind_Direction_NS | 1.30 | 0.086 | 15.17 | <0.001 | 1.13 | 1.47 |
| Post Arri | (Intercept) | 134.30 | <0.001 | 781680.50 | 0.00 | 134.30 | 134.30 |
| Post Arri | DBT | 0.119 | 0.104 | 1.13 | 0.257 | -0.086 | 0.323 |
| Post Arri | `MSLP (mbar)` | -0.131 | 0.003 | -47.69 | 0.00 | -0.137 | -0.126 |
| Post Arri | `Wind_Speed (m/s)` | -0.030 | 0.015 | -2.06 | 0.040 | -0.059 | -0.001 |
| Post Arri | Wind_Direction_EW | -0.555 | 0.099 | -5.61 | <0.001 | -0.748 | -0.361 |
| Post Arri | Wind_Direction_NS | 0.586 | 0.101 | 5.83 | <0.001 | 0.389 | 0.783 |

| **y.level** | **term** | **estimate** | **std.error** | **statistic** | **p.value** | **conf.low** | **conf.high** |
| --- | --- | --- | --- | --- | --- | --- | --- |
| Arri-1w | (Intercept) | -119.91 | <0.001 | -699365.00 | 0.00 | -119.91 | -119.91 |
| Arri-1w | DBT | -0.134 | 0.209 | -0.643 | 0.520 | -0.544 | 0.275 |
| Arri-1w | `MSLP (mbar)` | 0.123 | 0.006 | 21.21 | <0.001 | 0.112 | 0.135 |
| Arri-1w | `Wind_Speed (m/s)` | -0.035 | 0.048 | -0.729 | 0.466 | -0.130 | 0.059 |
| Arri-1w | Wind_Direction_EW | 1.42 | 0.163 | 8.66 | <0.001 | 1.10 | 1.74 |
| Arri-1w | Wind_Direction_NS | -1.07 | 0.150 | -7.14 | <0.001 | -1.37 | -0.780 |
| Arri-2w | (Intercept) | -82.86 | <0.001 | -311282.00 | 0.00 | -82.86 | -82.86 |
| Arri-2w | DBT | -0.483 | 0.210 | -2.30 | 0.022 | -0.896 | -0.071 |
| Arri-2w | `MSLP (mbar)` | 0.096 | 0.006 | 16.32 | <0.001 | 0.084 | 0.107 |
| Arri-2w | `Wind_Speed (m/s)` | 0.023 | 0.048 | 0.491 | 0.623 | -0.070 | 0.117 |
| Arri-2w | Wind_Direction_EW | 1.61 | 0.238 | 6.74 | <0.001 | 1.14 | 2.08 |
| Arri-2w | Wind_Direction_NS | -0.791 | 0.240 | -3.29 | <0.001 | -1.26 | -0.320 |
| Arri-1M | (Intercept) | 38.65 | <0.001 | 117382.20 | 0.00 | 38.65 | 38.65 |
| Arri-1M | DBT | -0.800 | 0.205 | -3.91 | <0.001 | -1.20 | -0.399 |
| Arri-1M | `MSLP (mbar)` | -0.014 | 0.006 | -2.45 | 0.014 | -0.025 | -0.003 |
| Arri-1M | `Wind_Speed (m/s)` | -0.023 | 0.046 | -0.504 | 0.614 | -0.114 | 0.067 |
| Arri-1M | Wind_Direction_EW | 1.12 | 0.193 | 5.80 | <0.001 | 0.742 | 1.50 |
| Arri-1M | Wind_Direction_NS | -0.287 | 0.188 | -1.53 | 0.127 | -0.656 | 0.082 |
| Arri-2M | (Intercept) | -81.32 | <0.001 | -231309.00 | 0.00 | -81.32 | -81.32 |
| Arri-2M | DBT | -1.09 | 0.206 | -5.31 | <0.001 | -1.50 | -0.691 |
| Arri-2M | `MSLP (mbar)` | 0.113 | 0.006 | 19.69 | <0.001 | 0.102 | 0.124 |
| Arri-2M | `Wind_Speed (m/s)` | -0.085 | 0.047 | -1.79 | 0.074 | -0.178 | 0.008 |
| Arri-2M | Wind_Direction_EW | 1.09 | 0.196 | 5.55 | <0.001 | 0.703 | 1.47 |
| Arri-2M | Wind_Direction_NS | -0.183 | 0.182 | -1.00 | 0.315 | -0.538 | 0.173 |
| Post Arri | (Intercept) | 258.67 | <0.001 | 1312513.00 | 0.00 | 258.67 | 258.67 |
| Post Arri | DBT | -0.077 | 0.200 | -0.387 | 0.699 | -0.469 | 0.314 |
| Post Arri | `MSLP (mbar)` | -0.250 | 0.006 | -44.63 | 0.00 | -0.261 | -0.239 |
| Post Arri | `Wind_Speed (m/s)` | -0.023 | 0.044 | -0.509 | 0.611 | -0.110 | 0.064 |
| Post Arri | Wind_Direction_EW | 0.168 | 0.193 | 0.871 | 0.384 | -0.210 | 0.545 |
| Post Arri | Wind_Direction_NS | -0.332 | 0.216 | -1.54 | 0.124 | -0.755 | 0.091 |

*Table S4b.* Multinomial logistic regression results for the coarse analysis -Rushikulya. Effect size is the log odds ratio, which is e^(estimate)

*Table S5a.* Average Area under the curve (AUC) of the receiver operating characteristic (ROC) curve for 1500 XGBoost models

| **Site** | **AUC** |
| --- | --- |
| Gahirmatha | 0.886 |
| Rushikulya | 0.949 |

*Table S5b.* Average XGBoost parameter importance for Gahirmatha (1500 models)

| **Feature** | **Gain** | **Cover** | **Frequency** |
| --- | --- | --- | --- |
| Wind Speed | 0.195888 | 0.203782 | 0.158398 |
| Lunar Phase | 0.182346 | 0.193106 | 0.158412 |
| Lunar State | 0.127584 | 0.14014 | 0.148771 |
| DBT | 0.113471 | 0.108259 | 0.140666 |
| MSLP | 0.108822 | 0.115076 | 0.156362 |
| Wind Direction NS | 0.101754 | 0.071891 | 0.0506 |
| Relative Humidity | 0.092775 | 0.097386 | 0.13651 |
| Wind Direction EW | 0.07736 | 0.07036 | 0.050281 |

*Table S5c.* Average XGBoost parameter importance for Rushikulya (1500 models)

| **Feature** | **Gain** | **Cover** | **Frequency** |
| --- | --- | --- | --- |
| DBT | 0.192079 | 0.162381 | 0.139737 |
| MSLP | 0.166729 | 0.150928 | 0.164987 |
| Relative Humidity | 0.157364 | 0.143972 | 0.129242 |
| Wind Direction EW | 0.112936 | 0.100613 | 0.065678 |
| Wind Direction NS | 0.110195 | 0.084468 | 0.072899 |
| Lunar Phase | 0.089654 | 0.128673 | 0.147027 |
| Lunar State | 0.089292 | 0.117103 | 0.149456 |
| Wind Speed | 0.081751 | 0.111862 | 0.130973 |

*Figure S1.* (a-b) El Nino Southern Oscilattion (ENSO) yearly maximum value across years with arribada, no arribada, and years preceding a year without an arribada for Gahirmatha and Rushikulya, respectively(c-d) Indian Ocean dipole (IOD) index yearly maximum value across years with arribada, no arribada, and years preceding a year without an arribada for Gahirmatha and Rushikulya, respectively

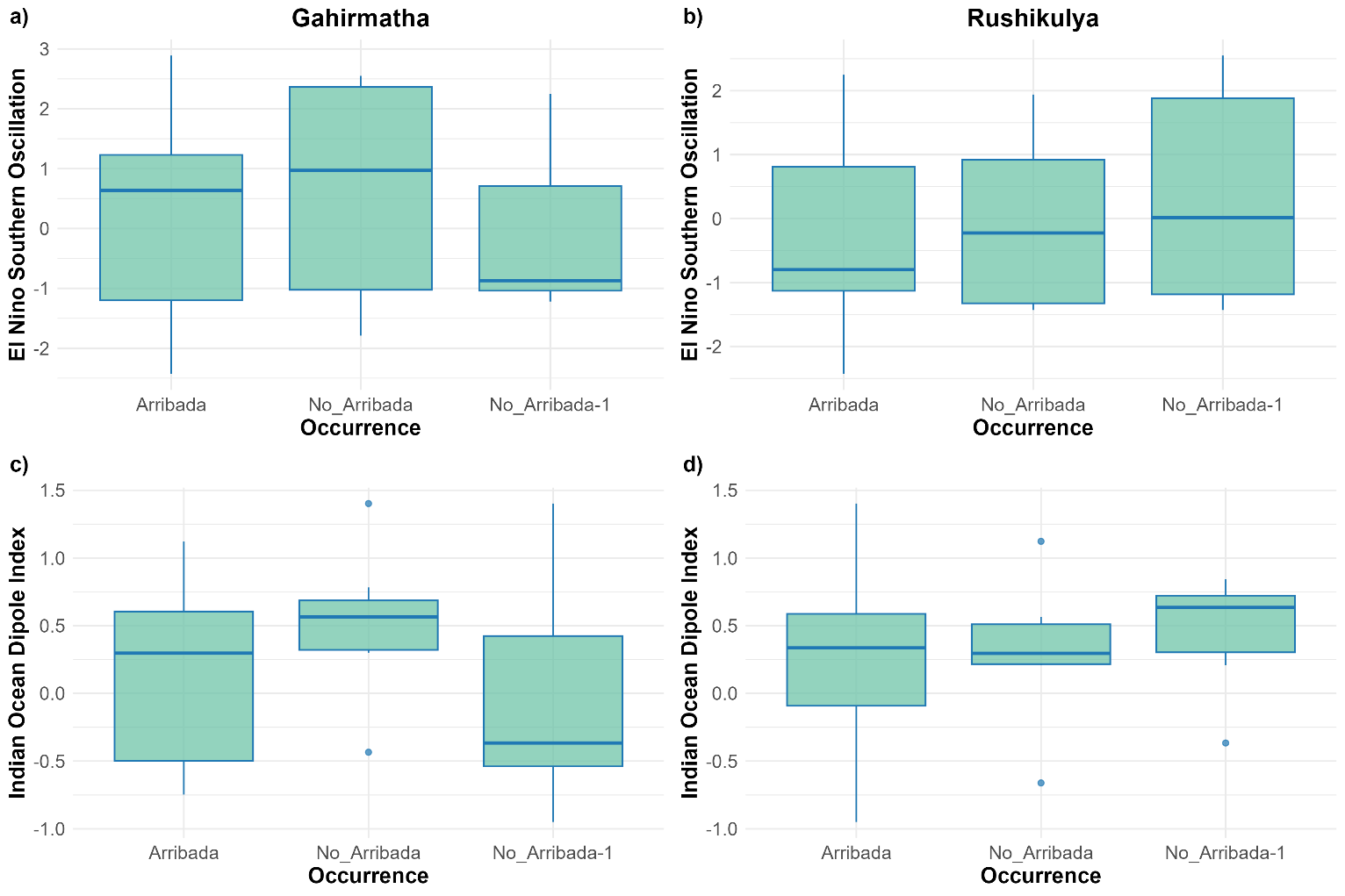

*Figure S2.* (a-b) Total rainfall for years with arribada, no arribada, and years preceding a year without an arribada for Gahirmatha and Rushikulya, respectively. (c-d) Mean sea level pressure (MSLP) across years with arribada, no arribada, years preceding a year without an arribada, and the arribada start date for Gahirmatha and Rushikulya, respectively

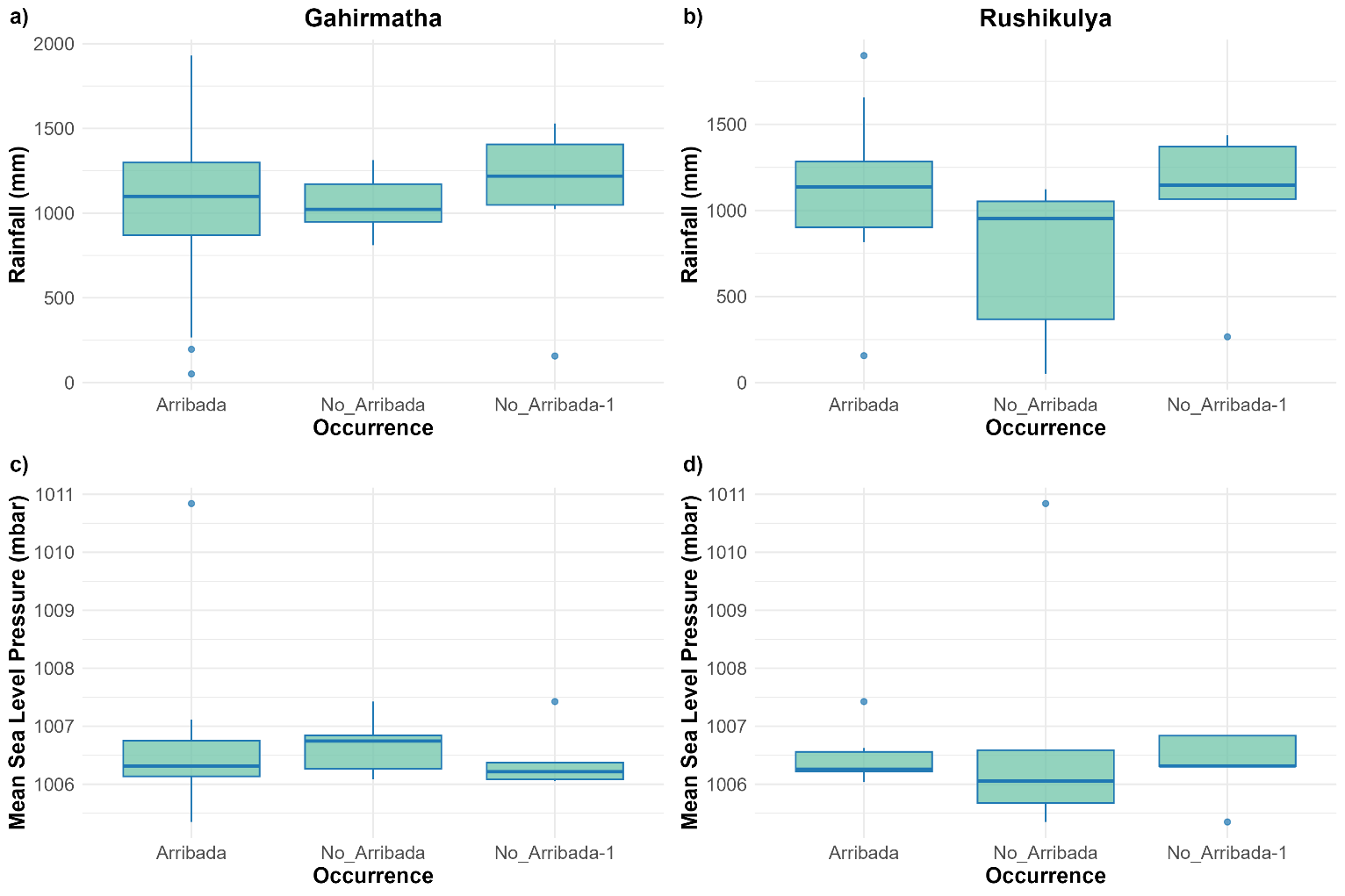

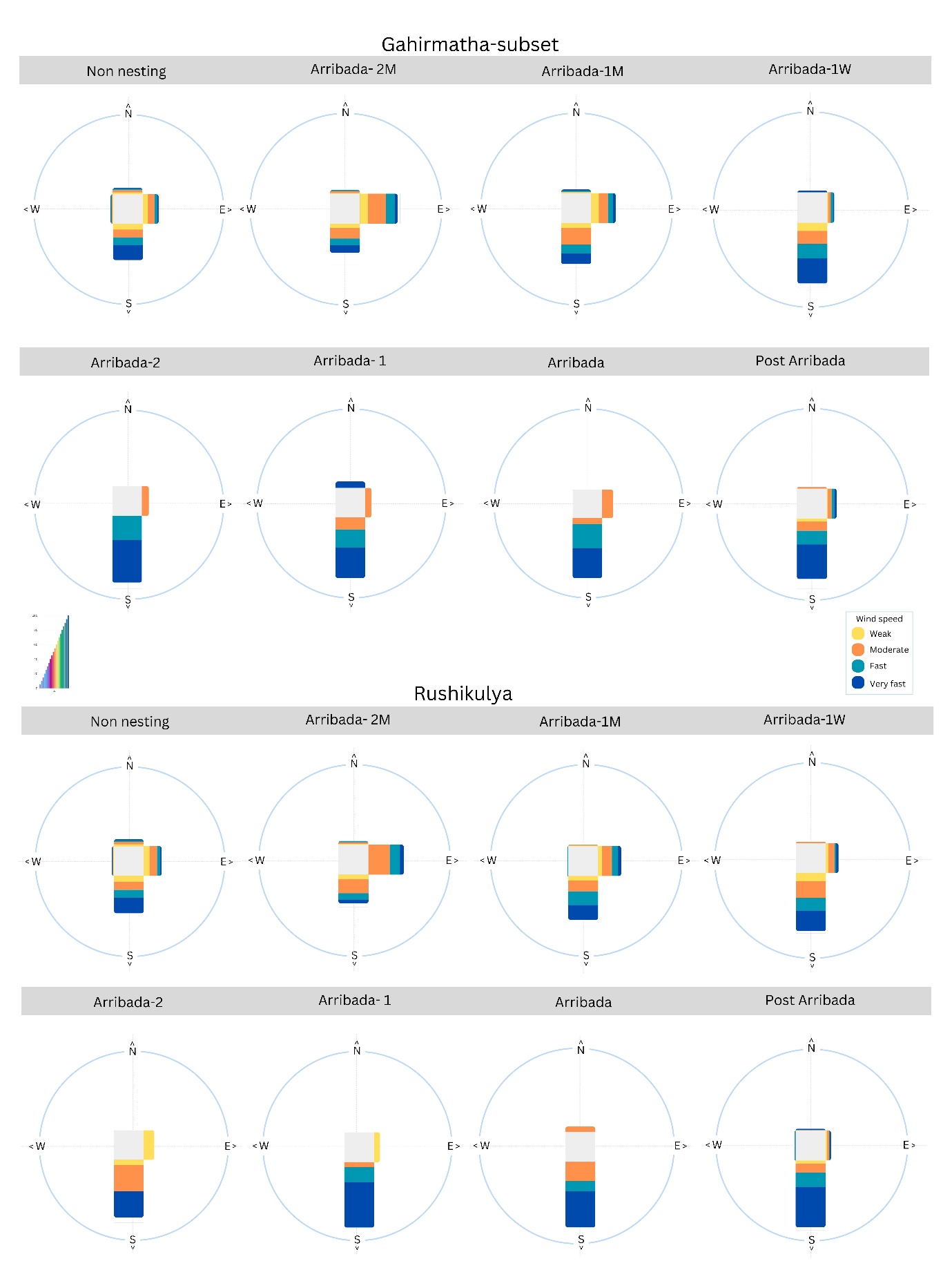
*Figure S3.* Wind speed and direction plots for a subset of Gahirmatha (1994-2019) and Rushikulya
